# Torsional Mechanics of Circular DNA

**DOI:** 10.1101/2024.10.08.617281

**Authors:** Gundeep Singh, Yifeng Hong, James T. Inman, James P. Sethna, Michelle D. Wang

## Abstract

Circular DNA found in the cell is actively regulated to an underwound state, with their superhelical density close to *σ* ∼ - 0.06. While this underwound state is essential to life, how it impacts the torsional mechanical properties of DNA is not fully understood. In this work, we performed simulations to understand the torsional mechanics of circular DNA and validated our results with single-molecule measurements and analytical theory. We found that the torque generated at *σ* ∼ - 0.06 is near but slightly below that required to melt DNA, significantly decreasing the energy barrier for proteins that interact with melted DNA. Furthermore, supercoiled circular DNA experiences force (tension) and torque that are equally distributed through the DNA contour. We have also extended a previous analytical framework to show how the plectonemic twist persistence length depends on the intrinsic bending persistence length and twist persistence length. Our work establishes a framework for understanding DNA supercoiling and torsional dynamics of circular DNA.

---

DNA topology plays a crucial role in genome organization and function, governing all processes involving DNA, including chromosome compaction, gene expression, DNA replication, and DNA repair. Notably, many chromosomes, such as mitochondrial DNA, bacterial chromosomes, and extrachromosomal DNA plasmids adopt circular forms. Because circular DNA lacks free ends for supercoiling dissipation, DNA supercoiling is constrained within the circular structure. Interestingly, these chromosomes maintain their supercoiling state despite alterations and perturbations by motor proteins, such as RNA polymerases and replisomes, or by the action of topoisomerases, which can relax or introduce DNA supercoiling. Maintenance of the topological integrity of these circular genomes is essential to their function.

Interestingly, mitochondrial DNA, bacterial chromosomes, and extrachromosomal plasmids are all maintained at a similar superhelical density of *σ* ∼ - 0.06 (where 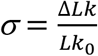 with *Lk*_0_ being the linking number of a relaxed DNA and Δ*Lk* being the change in the linking number due to supercoiling), despite their vast differences in genome sizes ^1^. The similarity in the supercoiling states suggests a universal need to maintain these genomes in a specific underwound state. However, due to our limited understanding of the torsional mechanical properties of circular DNA, it is difficult to assess how this supercoiling state might impact processes on DNA. It remains unclear how much torque this superhelical density generates in circular DNA and if such torque is sufficient to melt DNA or to stall motor proteins such as RNA polymerase during transcription. Answers to these questions require a thorough understanding of circular DNA torsional mechanics, which has been lacking thus far. Furthermore, plasmids have long been used as powerful tools to study DNA-protein interactions and DNA structure and function *in vitro* ^2^. Understanding the torsional mechanics of circular DNA could enhance our interpretative capabilities of plasmid-based biochemical investigations. There is also growing interest in supercoiled plasmids as potential substrates for gene therapy ^3^. Investigating the torsional stress of circular DNA with varying sizes and supercoiling densities could provide valuable insights into plasmid stability and efficient gene expression. This underscores the critical importance of accurately studying and measuring DNA torsional mechanics.

While the DNA torsional mechanics of linear DNA have been extensively studied theoretically ^4, 5^ and experimentally^6-15^, much less is known about that of circular DNA ^2, 16, 17^. We approach this problem by first employing coarse-grained Monte Carlo simulations to establish a framework to compute the torque in a circular DNA at a given supercoiling state, Δ*Lk* (Supplementary Material). In this approach, we discretize a circular DNA into a chain of smaller segments (Fig. 1a and b), impose an energetic cost of DNA bending (bending persistence length *A*) and twisting (twisting persistence length *C*), and allow the chain to equilibrate. The chain segments also interact with each other via electrostatic repulsion, modeled via a Debye-Huckel potential ^18, 19^.

**Figure 1:**
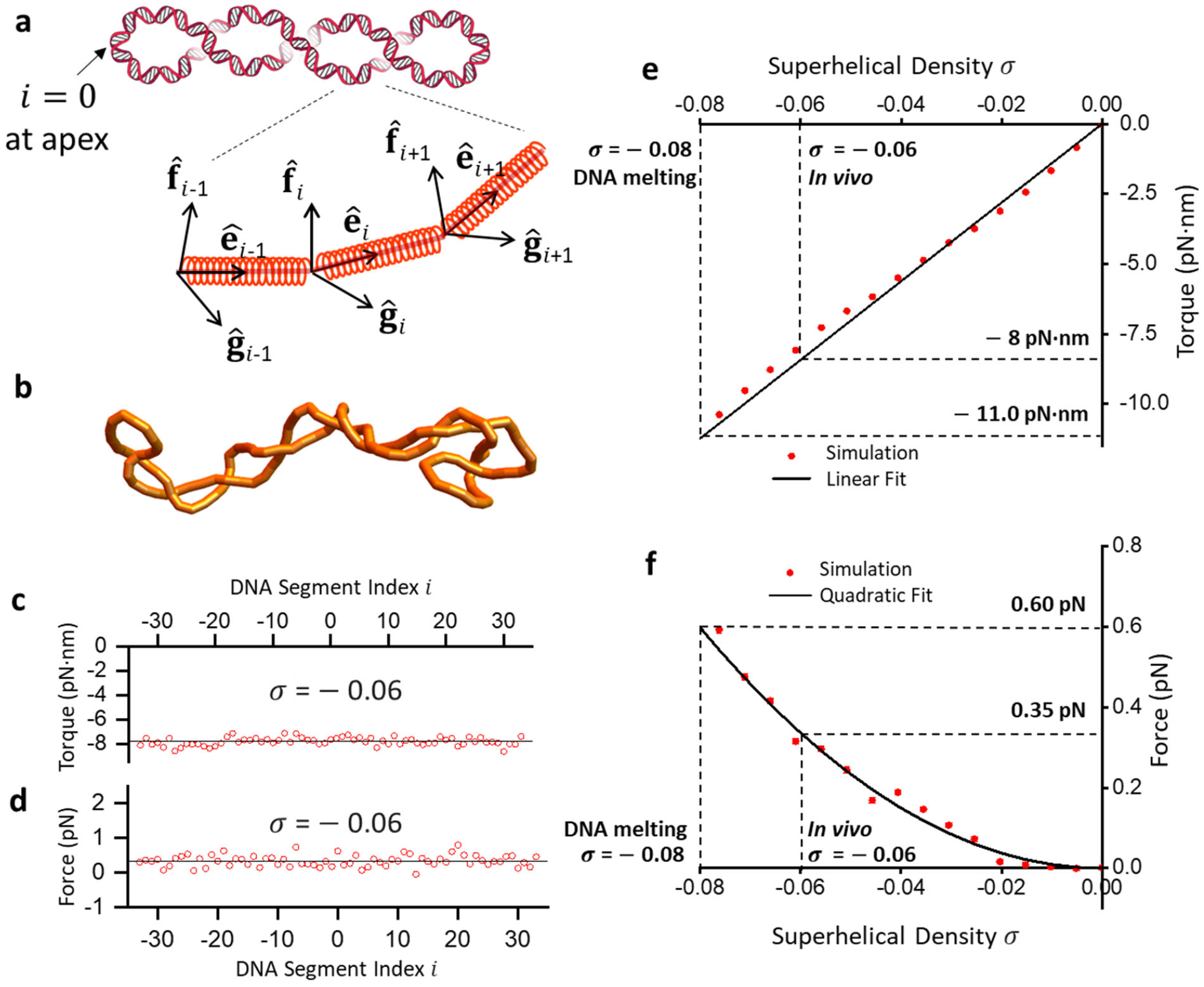
Simulating torsional mechanics of circular DNA. Monte Carlo simulations of circular DNA reveal the distribution and level of torque and force in a supercoiled circular DNA. (a) A cartoon illustrating the typical plectonemic configuration of supercoiled circular DNA. Equilibrium configurations of a supercoiled DNA are simulated using a discrete twistable worm-like chain model, where the orientation of each segment *i* is described using a set of orthonormal unit vectors 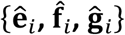 Here, 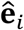 traces the DNA contour, while 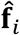 and 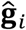 represent twist and are used to calculate the torque between segments. Each DNA segment is also assigned a spring along the segment with a large spring constant for force (tension) determination (see Supplementary Material) (b) Snapshot of a simulated configuration generated at *σ* = - 0.06. (c) and (d) The average torque and force for configurations with *σ* = - 0.06, plotted as a function of position along the contour. The DNA segment index is reordered such that *i* = 0 denotes the apex of the plectoneme for each configuration. e) Torque in the circular DNA increases with superhelical density and can be modeled well with a linear fit *τ* = *Pk*_B_*Tω*_0_*σ*, with *P* = 21 nm (see main text). (f) Force as a function of *σ* . A quadratic fit *F* = *a σ*^2^ results in the fit parameter *a* = 93.44 pN.

Because each segment has a twist coordinate, torque local to each segment can be obtained directly (Fig. 1c). We show that the torque remains the same for all chain segments, regardless of whether the segment is at the apex or the center of a plectoneme. Previous work on a filament with a uniform bending property shows that the torque and force (tension) along the filament should be constant when self-repulsion is neglected ^20^. Our results suggest that this remains valid even when self-repulsion is considered. This conclusion also extends to DNA with heterogeneous intrinsic parameters, where sequence-specific *A* and *C* vary across different regions (Fig. S1). In addition, torque *τ* increases linearly with *σ* (Fig. 1e). Since supercoiling of circular DNA induces plectoneme formation, the slope of the torque *τ* versus *σ* provides the plectonemic twist persistence length (*P*): *τ* = *Pk*_B_*Tω*_O_*σ*, where *k*_B_*T* is the thermal energy and 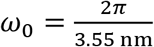 is a conversion factor related to the DNA helical rise. For *A* = 43 nm and *C*= 110 ^11^, the simulation yields *P* = 21 nm, close to what we have previously measured for a plectoneme in linear DNA ^11^. The agreement provides some validation for our simulation method.

There is often the perception that the force along a supercoiled circular DNA is nonexistent or so small that it is negligible. However, theoretical guidance on this has been lacking. Using the simulation to investigate tension in a circular DNA, we convert each segment along the DNA into a Hookean spring, which is used as a force readout (Fig. 1a) (Supplementary Material). Our results show that force indeed exists in supercoiled circular DNA (Fig. 1d). Importantly, the force is greater than zero, indicating that DNA is under tension when supercoiled. This force is present for each segment and remains the same for all chain segments. Therefore, the tension remains the same even when the DNA self-repulsion is considered. We further show that the force increases with an increase in the magnitude of *σ*, highlighting the presence of the tension as a direct consequence of the thermodynamic properties of supercoiled DNA (Fig. 1f).

These simulation results show that torque and force in a circular DNA increase with increased magnitude of superhelical density *σ*. At *σ* = -0.08, the torque reaches -11 pN·nm, which is required for the DNA melting transition ^9, 21^. Under this state, there is a concurrent force of 0.65 pN. At *σ* ∼ -0.06 of the mitochondrial DNA and bacterial chromosomes, the DNA experiences a torque of about -8 pN·nm, close to the DNA melting transition but slightly below the transition. Therefore, these circular chromosomes maintain a (-) supercoiling state close to that of DNA melting. Such (-) supercoiling could significantly reduce the energetic barrier for binding by various DNA-based processes, such as transcription factors and RNA polymerase^19, 22^, without melting DNA, which could create ssDNA that induces DNA damage responses. Although this torque is below what *E. coli* RNA polymerase can generate ^23, 24^, it could substantially impact RNA polymerase progression. Furthermore, at *σ* ∼ -0.06, the circular DNA has about 0.35 pN of force. This force is well below what could be generated by motor proteins, such as RNA polymerase ^25^, but is sufficiently large for impacting condensin and cohesin loop extrusion^26, 27^.

To validate the simulation results of circular DNA, we must be able to directly measure the torque and force required to supercoil circular DNA, which has not been experimentally feasible. On the other hand, direct torque and force measurements have been possible in single-molecule assays using an angular optical trap (AOT) where the torque in a linear DNA molecule is directly measured as the DNA is twisted to induce buckling under a constant force ^7, 11, 28-30^. Upon DNA buckling, a plectoneme is extruded with further supercoiling. The resulting plectoneme has a structure that resembles that of supercoiled circular DNA. If these two structures have the same mechanical properties, then it is possible to experimentally validate the simulation results of circular DNA using linear DNA. However, this assumption must first be rigorously examined.

We test this hypothesis by extending our computational framework to simulate the supercoiled linear DNA constrained to a fixed end-to-end extension (Supplementary Methods). Akin to our circular DNA simulations, we obtain the average torque and force as a function of supercoiling added to the DNA. We find that the magnitude of torque and force are uniformly distributed along the DNA contour regardless of whether a segment is in the extended region or the plectonemic region (Fig 2b). In Fig 2a, we plot the simulated torque and force versus turns partitioned into the plectonemic phase of the linear DNA and overlay the torque and force profiles of circular supercoiled DNA from Fig 1. The two conditions render nearly identical values. We also include theoretical predictions for torque and force of a purely plectonemic state of a linear DNA as given by the two-phase model by Marko^5^, which shows good agreement with our simulations. This confirms that one can alternatively study torsional mechanics of supercoiled circular DNA by obtaining the mechanics of a plectoneme formed on linear DNA.

**Figure 2:**
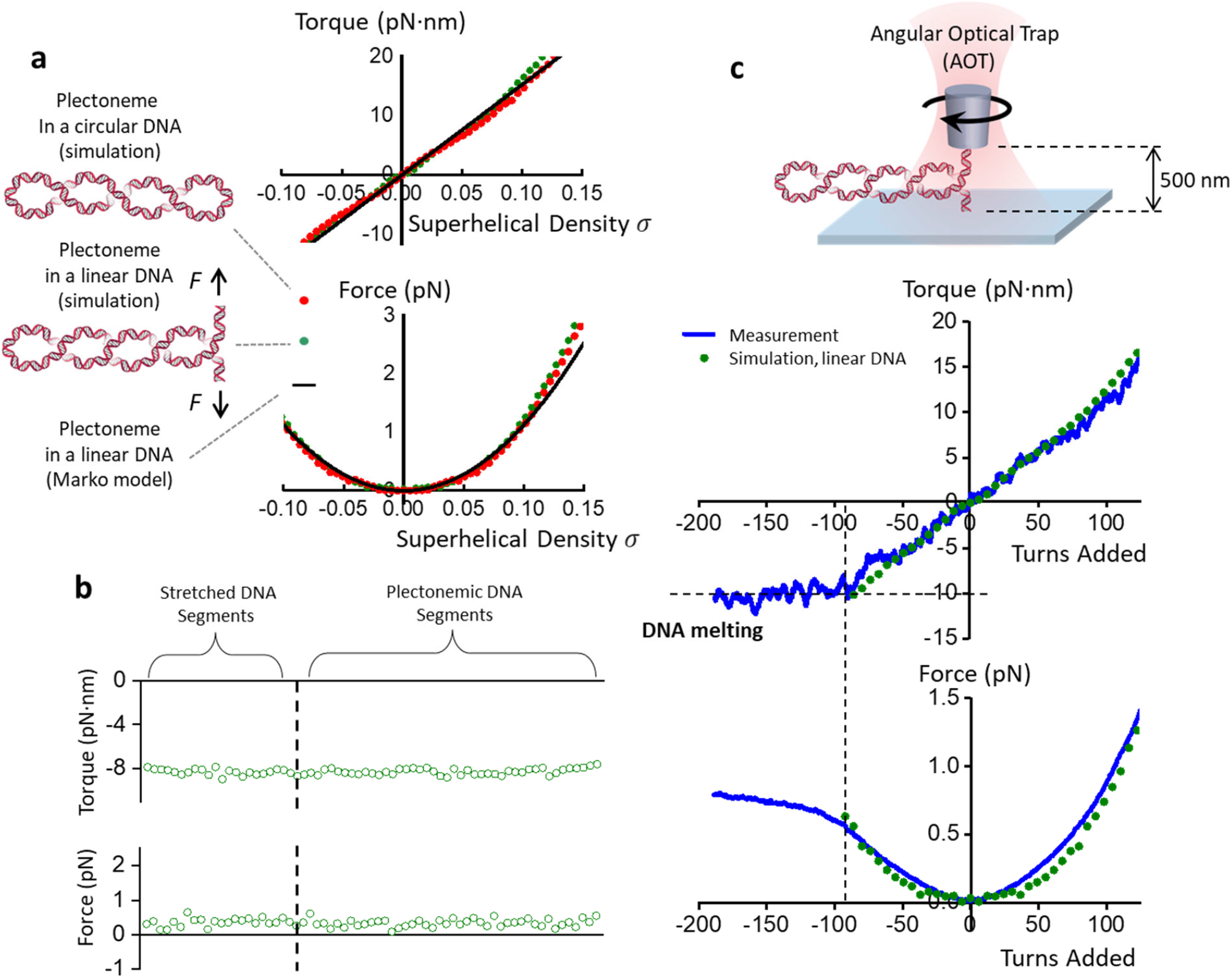
Simulating and comparing torsional mechanics of plectonemic linear DNA and circular DNA. (a) A comparison of simulated mechanical properties of a plectoneme formed on supercoiled linear DNA and circular DNA. Equilibrium configurations of linear DNA kept under a constant extension (1/9^th^ of total contour length) are simulated (see Supplementary Material) and the torque and force versus sigma partitioned into the plectoneme formed on the DNA are plotted. Torque and force profiles in the simulated circular DNA and linear plectonemic DNA are overlaid together as a function of superhelical density *σ*. Theoretical predictions of torque and force in a plectoneme of a linear DNA as given by the two-phase model of Marko^5^ are added for reference. (b) Average torque and force for linear DNA configurations with *σ* = - 0.06 analyzed by segment position, where each segment is categorized to be in either the extended or plectonemic regions. (c) Simulations of plectonemic linear DNA validated against single-molecule measurements. An angular optical trap (AOT) is used to experimentally measure torque and force of plectonemic DNA. A 12.7 kb linear DNA template is torsionally anchored between a coverslip and a nanofabricated quartz cylinder and maintained under a constant extension of 500 nm. Torque and force increase as turns are introduced to the DNA. In the negative supercoiling regime, the DNA undergoes a melting transition at around -90 turns. Simulations of linear DNA are overlaid with experimental measurements for comparison.

We follow this result and use the AOT to manipulate supercoiled linear DNA, directly measuring its torque and force. Briefly, in an AOT experiment, a linear DNA molecule has one end attached and torsionally constrained to a microscope coverglass, while the other end is attached and torsionally constrained to the bottom of a nanofabricated quartz cylinder^8, 11, 30-32^. A linearly polarized laser beam is used to exert force and torque on the cylinder which in turn exerts force and torque on the DNA (Fig. 2c). In these experiments, we use a linear DNA molecule of contour length 12.7 kb and fix its end-to-end extension to a relatively small magnitude of 500 nm (∼1/9th of the DNA contour length), similar to a method we previously developed ^11^. This configuration maximizes the torsional contribution of the plectonemic phase of the DNA while limiting interactions of the trapped cylinder with the coverglass surface.

Fig 2c shows the torque and force needed to supercoil a DNA kept at this small extension. When DNA is positively supercoiled, the torque rises linearly (indicating *P* = 22 nm), as shown in the circular DNA simulation. In the (-) supercoiling regime, the torque magnitude also increases linearly until *σ* ∼ -0.08, where DNA starts to undergo a melting transition, characterized by the plateauing of the torque data. Force rises almost quadratically in both the positive and negative regimes, although the DNA starts to melt when enough (-) supercoiling is built up (i.e., *σ* ∼ -0.08), after which the force nearly approaches a plateau. We find our simulated values of torque and force of linear DNA (in the B-DNA regimes) match with the experimental values very well before the melting transition. The AOT experiments, therefore, support and validate our simulated torsional parameters of circular DNA.

Our simulation results and experiments provide a value for the plectonemic torsional twist persistence length (*P*). However, *P* is not an intrinsic physical parameter. Instead, it is determined by two intrinsic parameters: *A*, the bending persistence length, and *C*, the twist persistence length. Although how *P* depends on *A* and *C* may be obtained numerically (Supplementary Fig. S2), there lacks an analytical expression for these dependencies, limiting an intuitive understanding of this parameter. Here, we show a method to derive a quasi-analytical expression by extending the work of Vologodskii et al. ^33^.

We start by considering a nicked circular DNA that can spontaneously form twist (Δ*Tw*) and writhe (Δ*Wr*). Then, the linking number (Δ*Lk*) will be given by Δ*Lk* = Δ*Tw* + Δ*Wr*. While the average of each of these three quantities would be zero, their thermodynamic fluctuations would be non-zero. In this case, it is assumed that variance in twist 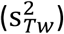 and variance in writhe 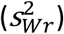 are independent of one another ^33, 34^, then the variance in 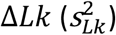 is given by:

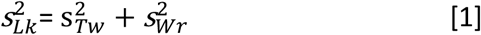

The twist persistence length (*P*) of circular DNA of contour length *L* is also related to the fluctuation of Δ*Lk* through the following equation ^35^:

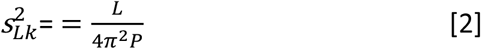

It is known that twist fluctuations only depend on *C* ^36^, while the writhe fluctuations for the circular DNA only depend on *A* ^37^.

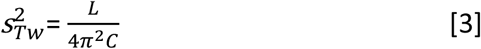

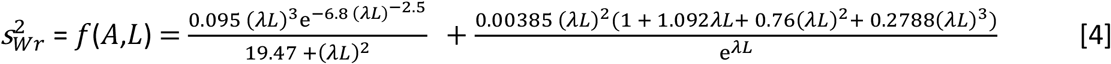

where, 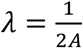 Here, Equation 4 is replicated from the empirical formula obtained by Shimada and Yamakawa ^37^. We validate the relations shown in equations 3 and 4 by simulating a nicked circular DNA and plotting the distributions for Δ*Tw* and Δ*Wr* (Fig. 3a; Supplementary Material). Substituting equations 2, 3, and 4 into equation 1, we get the following relationship:

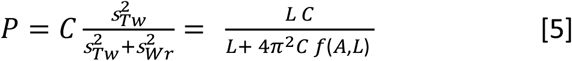

Equation 5 explicitly gives *P* as a function of DNA’s intrinsic parameters *A* and *C*. In Fig 3b, we validate equation 5 by varying *A* and *C* and comparing the simulated values of *P* against these theoretical estimates. We observe an excellent match between the two methods.

**Figure 3:**
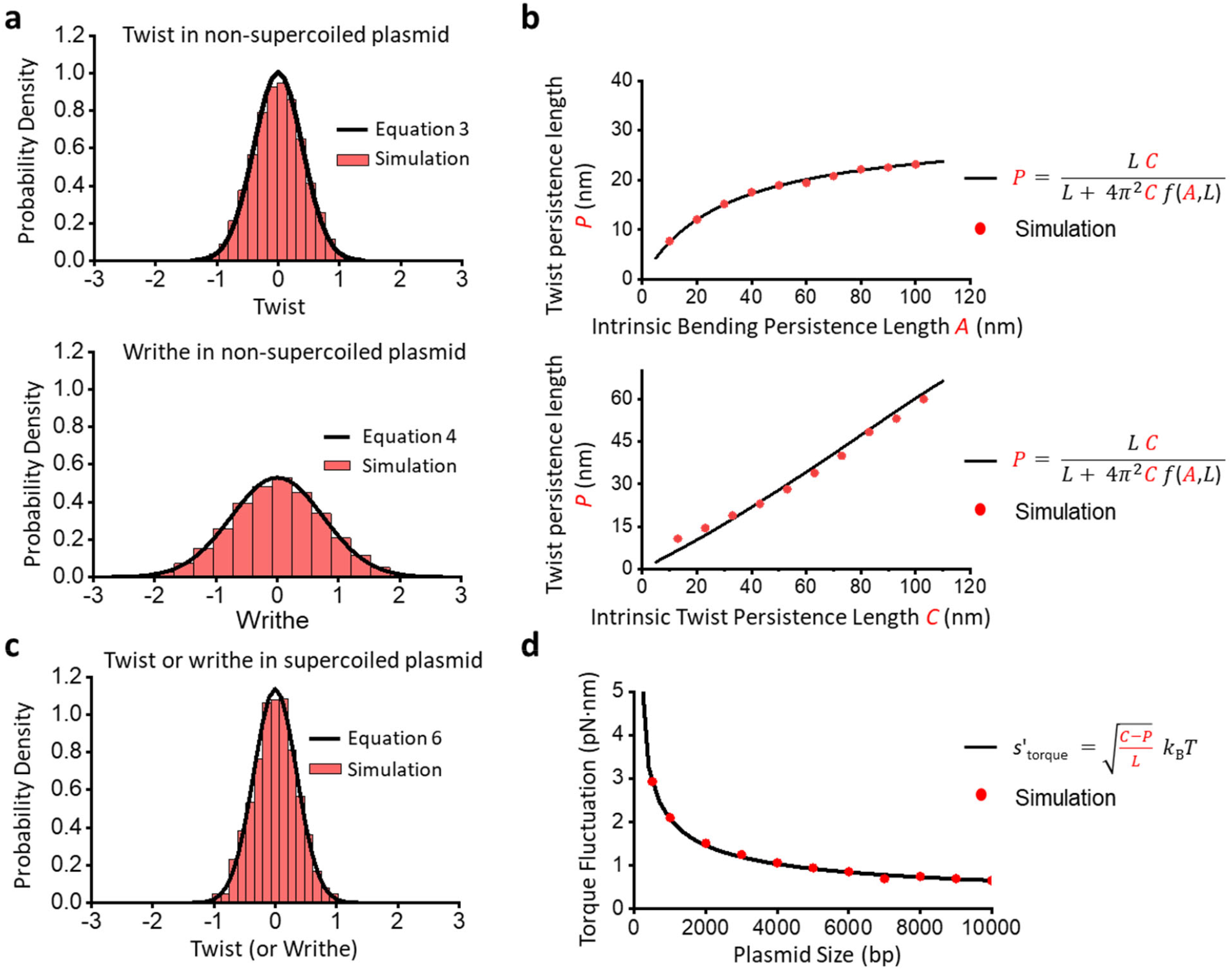
Circular DNA torque and its fluctuation are determined by the intrinsic parameters *A* and *C*. The plectonemic twist persistence length *P* depends on writhe and twist fluctuations in nicked DNA, given *A* and *C*. In supercoiled DNA, the torque fluctuations can be derived from these fluctuations. (a) A comparison between distributions of twist and writhe in nicked circular DNA as given by theory (Eqs. 3 and 4 in main text, respectively) and computed using simulations (see Supplementary Material) for *A* = 43 nm and *C*= 109 nm. (b) *P* is plotted as a function of *A* and *C*. In the top panel, *A* is varied with *C* fixed at 109 nm, and in bottom panel *C* is varied with *A* fixed at 43 nm. The closed form equation (Eq. 5) is compared to simulated values of *P* in both plots. (c) Twist (or writhe) distribution in a supercoiled circular DNA is shown as predicted by theory (Eq. 6) and computed through simulations at *σ* = 0 for *A* = 43 nm and *C*= 109 nm. (d) Torque fluctuations are plotted as a function of the size of the circular DNA using both theory (Eq. 7) and simulations at *σ* = -0.06 for *A* = 43 nm and *C*= 109 nm.

While torsional stiffness parameter *P* provides an excellent estimate for the average value of torque at a given supercoiling density (*τ* = *Pk*_B_*Tω*_O_*σ*), in a cellular system, the magnitude of torque would fluctuate due to the presence of thermal agitations. These fluctuations could further modulate the chances of local melting events in the circular DNA.

Here, we extend our fluctuation analysis to obtain the variance of torque in supercoiled circular DNA and how it relates to intrinsic parameters *A* and *C*. Contrary to the nicked DNA, where fluctuations in Δ*Tw* and Δ*Wr* are completely independent, in the case of a supercoiled circular DNA, the two fluctuations are fully coupled to one another. As Δ*Lk* is fixed, the relation Δ*Lk* = Δ*Tw* + Δ*Wr* allows us to obtain topological fluctuations in a supercoiled circular DNA (see Supplementary Material and ^38, 39^) The variance in the value of twist 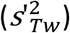 or writhe 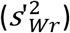 of a supercoiled circular DNA is then given by:

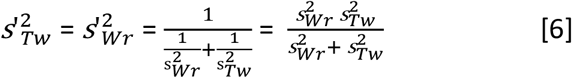

Converting the twist fluctuations to fluctuations in torque and substituting equations 3, 4, 5 into equation 6, we get:

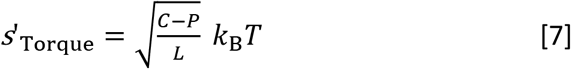

Fig 3c shows the comparison between torque distribution in a supercoiled circular DNA obtained from simulation against equation 7 for a given contour length of DNA (*σ* = 0), while Fig 3d shows how the torque fluctuations decrease with increasing DNA contour length at *σ* = -0.06. The simulated curve shows an excellent agreement with the theoretical estimate, thereby validating the results.

Our work establishes a framework for understanding DNA supercoiling and torsional dynamics of circular DNA using a combination of theoretical, computational, and experimental approaches. We show that a circular DNA at *σ* = -0.06 is under -8 pN·nm torque and 0.35 pN force, which could significantly impact processes on such a DNA substrate *in vivo*. Since small plasmids are also used in gene therapy and the FDA requires at least 70% of isoforms of plasmids to be supercoiled ^40-42^, torsional mechanics of a circular DNA may also play a role in the efficacy of plasmid-based gene therapy and help realize the full potential of this technology. We also demonstrate that circular DNA size significantly influences torque fluctuations, with smaller plasmids being more susceptible to melting phase transitions due to these fluctuations. These large fluctuations might not be preferred in plasmids *in vivo*, which need to finely regulate their transcription and replication levels. Hence, short plasmids (less than 1 kbp) might be evolutionarily disadvantageous and are rarely found in nature ^43, 44^. However, torque fluctuations in short plasmids could improve gene transfer efficiency in plasmid-based gene therapy. Therefore, in addition to topology, plasmid size is also a critical factor in selecting the optimal non-viral plasmid vector ^45^. Thus, understanding the torsional mechanics of circular DNA lays the crucial foundation for gaining insights into the fundamental processes of DNA and supporting practical applications.

We wish to thank Dr. C. Shonkwiler for their guidance in utilizing the conformal barycenter algorithm as part of the circular DNA simulations, Drs. J.F. Marko and B. Daniels for helpful discussions and constructive comments, and the Wang Lab for their useful feedback.

This work is supported by the National Institutes of Health grants R01GM136894 (to M.D.W.) and T32GM008267 (to M.D.W.). M.D.W. is a Howard Hughes Medical Institute investigator. The quartz cylinder fabrication was performed at the Cornell NanoScale Science & Technology Facility (CNF), a member of the National Nanotechnology Coordinated Infrastructure (NNCI), which is supported by NSF (NNCI-1542081).

