## Supplementary Materials for "Torsional Mechanics of Circular DNA"

### Methods

#### COARSE-GRAINED COMPUTER SIMULATIONS OF DNA

Simulation of the equilibrium conformations of DNA constructs are made with a Monte Carlo based method [1]. The DNA was modeled as a discrete twistable worm-like chain. Each segment $i$ of this chain has a segment length of $l_0$ and the positions of the endpoints (vertices) of the $i$ th segment are denoted by $\mathbf{r}_i = \{x_i, y_i, z_i\}^T$ and $\mathbf{r}_{i+1} = \{x_{i+1}, y_{i+1}, z_{i+1}\}^T$. A set of three orthonormal unit vectors $\{\hat{\mathbf{e}}_i, \hat{\mathbf{f}}_i, \hat{\mathbf{g}}_i\}$ are attached to each segment $i$ to keep track of structural and torsional deformations of the DNA as shown in Figure 1. Here, the vector $\hat{\mathbf{e}}_i$ describes the tangential component of the chain whose direction points along the segment $\mathbf{r}_{i+1} - \mathbf{r}_i$, while vectors , $\hat{\mathbf{f}}_i$ and $\hat{\mathbf{g}}_i$ store the twist information along the contour of the DNA. This model is also often referred to as the triad model [2, 3]. Through the course of the simulation, DNA conformations are explored by directly transforming $\hat{\mathbf{e}}_i$ and $\hat{\mathbf{f}}_i$ and the orthogonality is maintained by fixing $\hat{\mathbf{g}}_i = \hat{\mathbf{e}}_i \times \hat{\mathbf{f}}_i$. In the following subsections we first describe the energy terms for each generated configuration of the DNA chain and then elaborate on the Monte Carlo procedure we have employed to sample equilibrium configurations.

##### Energy terms of the model

To model the DNA, the energy required to bend and twist the DNA is considered as well as the electrostatic interaction from the charged DNA backbone. Additionally, to detect force in the DNA, segments are allowed to vary their lengths away from $l_0$, which results in a harmonic penalty term modeled by a Hookean spring elastic energy. Therefore, for any given DNA configuration, its energy consists of four main terms: bending energy ($E_{\mathrm{b}}$), torsional energy ($E_{\mathrm{t}}$), electrostatic repulsion energy ($E_{\mathrm{r}}$), and the elastic spring energy ($E_{\mathrm{s}}$). The bending energy ($E_{\mathrm{b}}$) gets a contribution from each consecutive pair of segments and depends on the bending angle $\beta_i = \cos^{-1}(\hat{\mathbf{e}}_i \cdot \hat{\mathbf{e}}_{i+1})$. For circular DNA simulations, the index $i$ spans from 1 to $N$, where $N$ is the total number of segments and $\beta_N$ is the angle between the first and last segment. For linear DNA simulation, $i$ spans from 1 to $N$-1 , since the first and last segments are no longer connected. The total bending energy is therefore given by:

$$E_b = \frac{\alpha}{2} k_B T \sum \beta_i^2 \qquad (1)$$

Here, $k_B T$ is the thermal energy unit and the pre-factor $\alpha$ is the dimensionless bending rigidity constant estimated by solving the following equation [4]:

$$\frac{2A}{l_0} = \frac{1 + \langle \cos\theta \rangle}{1 - \langle \cos\theta \rangle}$$

where, $A$ is the bending persistence length (Table S1) and

$$\langle \cos\theta \rangle = \frac{\int_0^{\pi} d\theta \cos\theta \, \sin\theta \, \exp(-\alpha\theta^2/2)}{\int_0^{\pi} d\theta \, \sin\theta \, \exp(-\alpha\theta^2/2)}$$

Similar to Eq. 1, the contribution to torsional energy is given by,

$$E_t = \frac{C k_B T}{2 l_0} \sum \psi_i^2 \quad (2)$$

Here, $C$ is DNA's intrinsic twist persistence length (Table S1) and $\psi_i$ is the local twist angle between segments $i$ and $i+1$ and is given by [5]:

$$\psi_i = \tan^{-1}\left(\frac{\hat{\mathbf{f}}_{i+1} \cdot \hat{\mathbf{g}}_i - \hat{\mathbf{g}}_{i+1} \cdot \hat{\mathbf{f}}_i}{\hat{\mathbf{f}}_{i+1} \cdot \hat{\mathbf{f}}_i + \hat{\mathbf{g}}_{i+1} \cdot \hat{\mathbf{g}}_i}\right) \quad (3)$$

Next, we estimate the electrostatic repulsion energy from the charged DNA backbone in an aqueous solution of salt using the Debye-Huckel approximation [6]. Here, the electrostatic contribution due to pairs of all non-consecutive segments of DNA are considered:

$$E_r = k_B T \xi^2 l_b \sum_{i=1}^{N-2} \sum_{j=i+2}^{N} E_r^{ij} \qquad (4)$$

where $\xi$ is the effective linear charge density (Table S1) of the DNA segment, $l_b$ is the Bjerrum length in water at temperature $T$, and $E_r^{ij}$ is computed by solving the following double integral numerically:

$$E_r^{ij} = \int_0^{l_0} d\lambda_i \int_0^{l_0} d\lambda_j \frac{\exp(-\kappa r_{ij})}{r_{ij}} \quad (5)$$

Here, the integration is done along the two segments; $\lambda_i$ and $\lambda_j$ are the distance parameterization along the segments, $r_{ij}$ is the distance between the two positions on the segments corresponding to the integration parameters $\lambda_i$ and $\lambda_j$ correspond, and finally, $\kappa$ is the inverse of the Debye length (Table S1). To save computational time, a tabulation of the double integral (Eq. 5) was used by creating a look-up table for a range of 3D segment orientations as described by Klenin and Langowski [7]. During the simulation, a multivariate linear interpolation is used to estimate $E_r^{ij}$ for each pair of $i$ and $j$ as described by Eq. 5.

Finally, the chain segments are allowed to vary their lengths away from the equilibrium length $l_0$. The variation in segment lengths is then used to detect force in the DNA. This results in a harmonic penalty term modelled using a Hookean elastic energy:

$$E_s = \frac{k}{2} \sum_{i=1}^{N} (l_0 - l_i)^2$$

Here, $l_i$ is the length of segment $i$ (magnitude of the vector difference $\mathbf{r_{i+1}}$ - $\mathbf{r_i}$) and $k$ is the spring constant associated with length variation of the segments. The spring constant is selected to be sufficiently large so that the extension variation remains much smaller than $l_0$ ensuring it does not interfere with the DNA configuration landscape. However, it is important to note that the variance in force increases linearly with $k$. Therefore, $k$ cannot be so high that the estimated force fails to converge within a practical computational time frame.

**Simulation procedure**

1. Circular DNA Simulation

We simulate equilibrium configurations of DNA under different values of supercoiling $\Delta Lk_0$ using the Metropolis Monte Carlo procedure [1]. We start each simulation with a circular chain with all the turns partitioned into twist. The configuration landscape is explored by applying three different kinds of structural moves: crankshaft, twist rotation, and segment length variation. Each move is selected with equal probability, and each Monte Carlo step involves a random choice of one of these moves. Crankshaft rotations sample DNA contour geometry while keeping the end-to-end extension fixed by rotating a segment of the chain about a line that passes through its endpoints [1]. We sampled the degree of crankshaft rotations from a uniform distribution between [−30°,30°]. Twist rotations vary twist angle $\psi_i$ between randomly selected segments $i$ and $i+1$, with angles sampled from a uniform distribution between [−50°,50°]. Finally, segment length variation moves increase or decrease the length $l$ of a randomly chosen force detector spring segment by a random value drawn from a uniform distribution of [-1.5 nm, 1.5 nm]. This move changes the end-to-end extension of the chain away from zero. Therefore, after every segment length variation move, we close the open polygon by computing the conformal barycenter of the configuration and transforming the conformation as detailed in the algorithm developed by Cantarella and Schumacher [8].

Each new configuration generated is checked for three conditions before it is accepted. First is the Metropolis-Hasting criterion with the energy $E$ = $E_{\mathrm{b}}$ +$E_{\mathrm{t}}$ + $E_{\mathrm{r}}$ + $E_{\mathrm{s}}$. Second, we ensure that the linking number remains close to the turns added to the DNA $\Delta Lk_0$ (within the tolerance of 0.05 turns). This is done by computing $\Delta Lk$ = $\Delta Tw$ + $\Delta Wr$ by directly estimating twist ($\Delta Tw$ = $\Sigma\psi_i$) and writhe (see Klenin and Langowski [9]). This condition is omitted in the nicked circular DNA simulation shown in Figure 3. Finally, it is required that the chain remains unknotted through the course of the simulation. This is evaluated by computing the Alexander Polynomial of the chain and if a trial move results in a knotted topology, it is rejected [10]. The overall acceptance rate for the step sizes described is between 30% and 50% across all superhelical densities explored. Every 500$^{\mathrm{th}}$ accepted configuration is stored and analyzed for torque and force measurements.

2. Linear DNA Simulation

We have also simulated linear DNA under constant extension to mimic the AOT experimental setup (Fig 2). While many details of the linear DNA simulation remain the same as the circular DNA simulation, in this sub-section, we have highlighted the differences between the two simulations. In the linear DNA simulations, we start from a completely straight chain with all its turns partitioned into twist. We then apply the three structural moves as described above (crankshaft, twist rotations, and segment length variation). To maintain a constant extension in this case, we can no longer use conformal barycenter. Hence, we simply use a stiff spring with no equilibrium length to anchor the chain to an end point $Z_0$ distance away from the origin. An energy term ($\frac{k\prime}{2}\ (Z_0 - Z)^2$) is added to the energy equation with $k'$ being the stiff spring constant and Z being the extension of the current configuration. The configuration is again checked for the three conditions before it is accepted: Metropolis-Hasting criterion, linking number conservation, and unknot topology check. Additionally, we have added strict hard wall boundaries to mimic the AOT setup at the surface and the quartz cylinder, where a configuration is rejected if any of its segments crosses those two hard wall surfaces.

**Configuration Analysis**

1. Plectoneme Apex Identification in Circular DNA

Before computing the torque and force of saved equilibrium configurations, we re-numbered our segments based on their position. For the circular DNA simulation, we have used an algorithm proposed by Vologodskii et al. [1] where the maxima in the local writhe contribution of the configuration are used to identify plectonemic apex positions. Here, we have used the apex with the largest local writhe contribution to be our index position 0. The remaining indexes are circularly shifted around the 0-apex position.

2. Separating Plectoneme and Extended DNA Contribution in Linear DNA

Saved linear DNA configurations are analyzed to divide their segments into extended and plectonemic phases. Here, the beginning and end points are identified for each plectonemic domain formed by calculating the longest pair of segment indexes (in contour length) with their distances (in extension) less than 10 nm. The segments outside of the domain are automatically categorized as extended segments. The local torque and force of the two phases are grouped together for comparison in Fig 2b.

#### DERIVING TORQUE FLUCTUATIONS IN A SUPERCOILED CIRCULAR DNA

While twist and writhe are independent in a nicked circular DNA, they become fully interdependent in a supercoiled circular DNA due to the presence of topological constraint. Since a supercoiled circular DNA lacks free ends for supercoiling dissipation, the linking number $\Delta Lk$ remains fixed in a supercoiled circular DNA. The linking number is the sum of the twist and writhe $\Delta Lk = \Delta Tw + \Delta Wr$ and therefore with a fixed $\Delta Lk$ any change in $\Delta Tw$ must be exactly balanced by an opposite change in $\Delta Wr$. This means that the fluctuations (or the variances) of $\Delta Tw$ and $\Delta Wr$ are the same because both quantities respond equally, but in opposite directions, to maintain the constant sum. In this derivation, we extend the framework by Levens and Crothers [11] and use the direct coupling of $\Delta Tw$ and $\Delta Wr$ in a supercoiled circular DNA to derive how twist (and by conjunction, torque) fluctuates in a supercoiled circular DNA.

Given a fixed linking number $\Delta Lk = \lambda$, the probability distribution of observing the twist $\Delta Tw = x$ follows the conditional distribution $P_{\Delta Tw\,|\Delta Lk}\,(x|\lambda)$, since $\Delta Tw$ depends on the fixed value of $\Delta Lk$. Using the Bayes' rule for continuous variables, we can write this distribution as:

$$P_{\Delta Tw\,|\Delta Lk}\,(x|\lambda) = \frac{P_{\Delta Tw}\,(x)\;P_{\Delta Lk\,|\Delta Tw}\,(\lambda|x)}{P_{\Delta Lk}\,(\lambda)}$$

where,

- $P_{\Delta Tw}(x)$ is the probability density of unconstrained twist, which describes how twist fluctuates independently;
- $P_{\Delta Lk\,|\Delta Tw}\,(\lambda|x)$ is the conditional probability of observing $\Delta Lk = \lambda$ given $\Delta Tw = x$. We use the coupling of $\Delta Tw$ and $\Delta Wr$ here to rewrite $P_{\Delta Lk\,|\Delta Tw}\,(\lambda|x) = P_{\Delta Wr}\,(\lambda - x)$, where the latter is the probability density of unconstrained writhe at $\Delta Wr = \lambda - x$; and
- $P_{\Delta Lk}\,(\lambda)$ is the total probability of observing $\Delta Lk{=}\lambda$ which normalizes the conditional distribution $P_{\Delta Tw\,|\Delta Lk}$.

In this derivation, we assume that $P_{\Delta Tw}(x)$ and $P_{\Delta Wr}(\lambda - x)$ both follow Gaussian distributions, which is well supported by our simulations (Fig. 3a) and previous literature [11]. Then $P_{\Delta Tw}(x)$ and $P_{\Delta Wr}(\lambda - x)$ are given by:

$$P_{\Delta Tw}(x) = \frac{1}{\sqrt{2\pi\, \mathrm{s}_{Tw}^2}} \exp(-\frac{(x - \langle\Delta Tw\rangle)^2}{2\mathrm{s}_{Tw}^2})$$

$$P_{\Delta Wr}(\lambda - x) = \frac{1}{\sqrt{2\pi\, \mathrm{s}_{Wr}^2}} \exp(-\frac{((\lambda - x) - \langle\Delta Wr\rangle)^2}{2\mathrm{s}_{Wr}^2})$$

where, $\mathrm{s}_{Tw}^2$ and $\mathrm{s}_{Wr}^2$ are the unconstrained DNA's twist and writhe variance as given by Eqs. 3 and 4 respectively in the main text and $\langle\Delta Tw\rangle$ and $\langle\Delta Wr\rangle$ are the average values of twist-writhe partitioning at the fixed $\Delta Lk$. Furthermore, due to linking number conservation equation, we can re-write $\big(((\lambda - x) - \langle\Delta Wr\rangle\big) = (\langle\Delta Tw\rangle - x)$. Therefore, $P_{\Delta Wr}(\lambda - x)$ becomes:

$$P_{\Delta Wr}(\lambda - x) = \frac{1}{\sqrt{2\pi\, s_{Wr}^2}} \exp(-\frac{(x - \langle \Delta Tw \rangle)^2}{2 s_{Wr}^2})$$

Finally, $P_{\Delta Lk}\ (\lambda)$ normalizes the conditional distribution and is the total probability of observing $\Delta Lk = \lambda$, given by:

$$P_{\Delta Lk}\ (\lambda) = \int_{-\infty}^{\infty} dx\ P_{\Delta Tw}\ (x)\ \ P_{\Delta Lk\ |\Delta Tw}\ (\lambda | x)$$

We denote $P_{\Delta Lk}\ (\lambda)$ as ${}^{1}/_{\mathrm{P}_0}$ for simplicity and will replace $\mathrm{P}_0$ such that $P_{\Delta Tw\ |\Delta Lk}\ (x|\lambda)$ is normalized.

Substituting above expressions into the equation for $P_{\Delta Tw\ |\Delta Lk}\ (x|\lambda)$ we get:

$$P_{\Delta Tw\ |\Delta Lk}\ (x|\lambda) = P_0\ \frac{1}{2\pi\, s_{Tw}\, s_{Wr}}\ \exp(-\frac{(x - \langle \Delta Tw \rangle)^2}{2} (\frac{1}{s_{Tw}^2} + \frac{1}{s_{Wr}^2})\ )$$

Notice that above distribution results in another Gaussian distribution, with its variance $s'^2_{Tw}$ related to the variance of unconstrained twist and writhe ($s_{Tw}^2$ and $s_{Wr}^2$, respectively) by the relation

$$s'^2_{Tw} = \frac{1}{\frac{1}{s_{Wr}^2} + \frac{1}{s_{Tw}^2}} = \frac{s_{Tw}^2 s_{Wr}^2}{s_{Tw}^2 + s_{Wr}^2}$$

The prime symbol is used to differentiate fluctuations in supercoiled DNA from those in unconstrained DNA. Substituting $s'^2_{Tw}$ and normalizing the distribution, $P_{\Delta Tw\ |\Delta Lk}\ (x|\lambda)$ becomes:

$$P_{\Delta Tw\ |\Delta Lk}\ (x|\lambda) = \frac{1}{\sqrt{2\pi\, s'^2_{Tw}}} \exp(-\frac{(x - \langle \Delta Tw \rangle)^2}{s'^2_{Tw}})$$

We can further express $s'^2_{Tw}$ in terms of the torsional parameters of the circular DNA, $C$ and $P$, using the relations derived in the main text. Rearranging Eq. 5 (main text), we get:

$\frac{s^2_{Tw}}{s^2_{Tw}+s^2_{Wr}} = \frac{P}{C}$ and $s^2_{Wr} = \frac{L}{4\pi^2} \left(\frac{C-P}{P\,C}\right)$

Substituting above equations, we get the following relation for $s'^2_{Tw}$:

$$s'^2_{Tw} = \frac{L}{4\pi^2} \left(\frac{C-P}{C^2}\right)$$

Once we know the twist fluctuation, we can then finally describe how torque fluctuation depends on DNA's torsional persistence lengths ($C$ and $P$):

$$s'^2_{\text{Torque}} = \left(\frac{2\pi\, k_{\text{B}}T\, C}{L}\right)^2 \frac{L}{4\pi^2} \left(\frac{C-P}{C^2}\right) = \frac{C-P}{L} \left(k_{\text{B}}T\right)^2$$

The above expression is plotted in Fig 3d of the main text as a function of different contour lengths $L$ of the DNA.

**ANGULAR OPTICAL TWEEZERS WITH CONSTANT EXTENSION**

The torque measurements were performed on the angular optical trap (AOT), which can simultaneously apply rotation and measure force, extension, and torque of a biomolecule[12-14]. In this work, a 12,688 bp DNA template, prepared as previously described[14-19], was torsionally constrained between a streptavidin-coated (Agilent, SA-10) coverslip surface and an anti-digoxigenin-coated (Roche, 11333089001) birefringent quartz cylinder. The cylinder was held at a constant extension of 500 nm and rotated at 2 turns/s to measure the twist persistence length of a plectonemic DNA as previously described[18]. The measured force was smoothed by a sliding window of 0.5 s with its offset removed by the theoretical modified Marko-Siggia model

[20]. The measured torque was smoothed by a sliding window of 2s. The measurements were carried out in a previously described topoisomerase reaction buffer (10 mM Tris-HCl pH 8.0, 50 mM KCl, 50 mM NaCl, 3 mM $MgCl_2$, 1 mM ATP, 0.1 mM EDTA, and 1.5 mg/mL β-casein (Sigma, C6905) at room temperature (23 $^{o}$C).

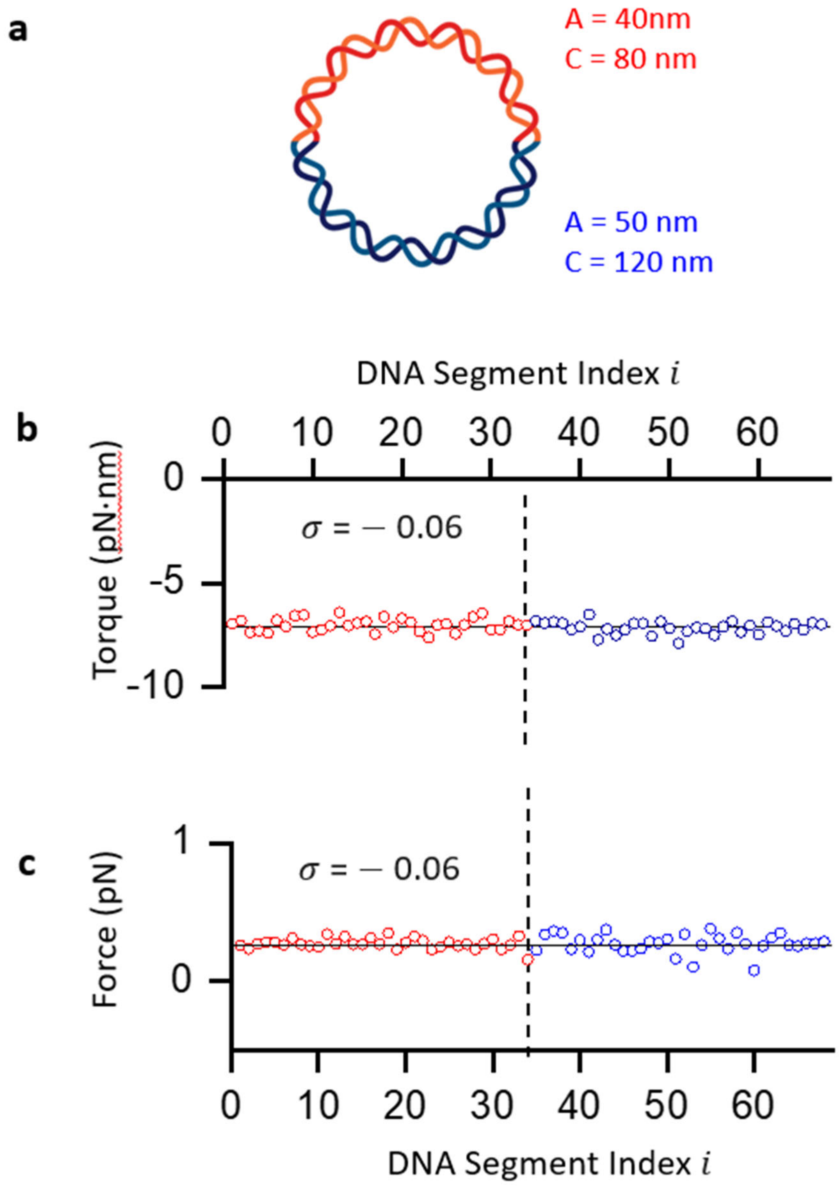


**Figure S1**: **Distribution of torque and force in a heterogeneous DNA template.** (a) Circular DNA is divided into two regions, each defined by different values of $A$ and $C$ based on sequence-dependent variations measured by Skoruppa et al.[3] Simulations were performed with $A$ = 40 nm and $C$ = 80 nm assigned for half of the DNA region and $A$ = 50 nm and $C$ = 120 nm assigned for the other half of the DNA contour. (b) and (c) The average torque and force of are plotted as a function of DNA segment index. The data are colored for the two regions based on the distinct intrinsic parameters (red for $A$ = 40 nm and $C$ = 80 nm and blue for $A$ = 50 nm and $C$ = 120).

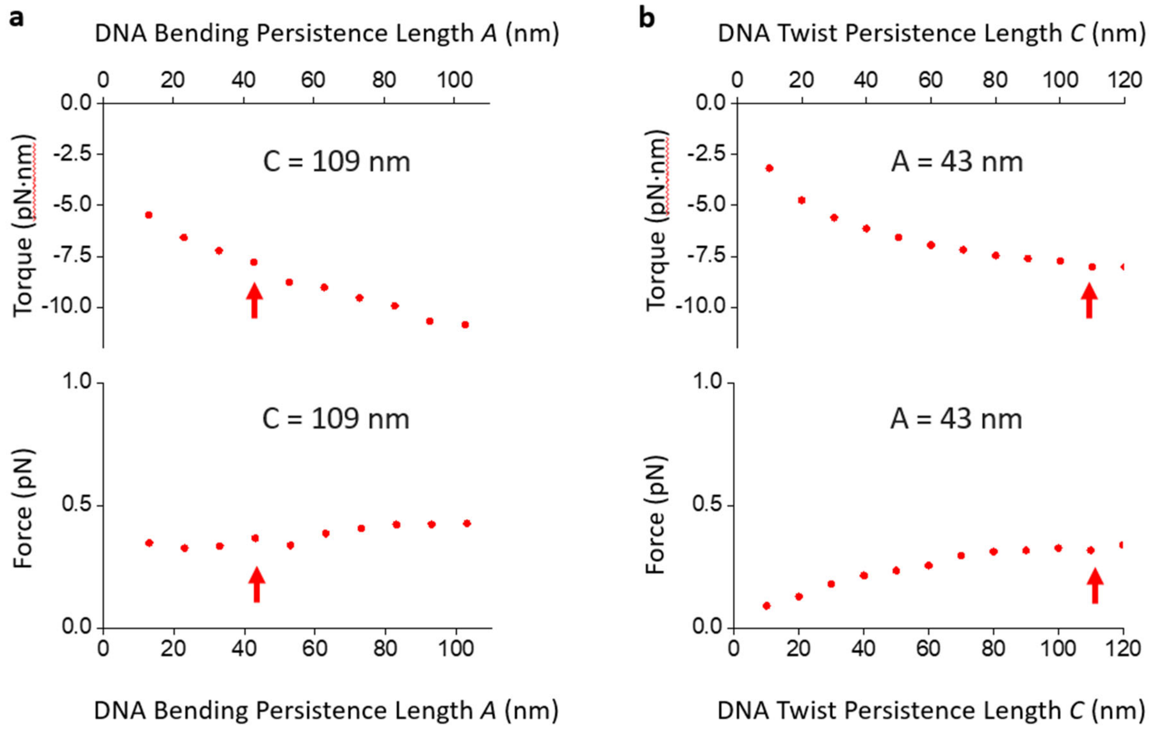


**Figure S2: Effect of DNA bending and twist persistence lengths on torque and force at $\sigma$ = - 0.06.** (a) Average torque and force generated from simulations of circular DNA with varying bending persistence length $A$, while twist persistence length $C$ is fixed at 109 nm. Red arrows highlight values at $A$ = 43 nm. (b) Average torque and force simulated with varying $C$, while $A$ is fixed at 43 nm. Red arrows highlight values at $C$ = 109 nm

| Parameter | Value |
|---|---|
| DNA length | 2000 bp |
| Segment length ($l_0$) | 10 nm |
| Temperature ($T$) | 297 K |
| Linear persistence length ($A$) | 43 nm[21] |
| Twist persistence length ($C$) | 109 nm[14, 15, 18] |
| Spring Constant ($k$) | 10 pN/nm |
| Inverse Debye Length ($\kappa$) | 1.35 $nm^{-1}$ [6] |
| Effective Linear Charge Density ($\xi$) | 5.74 e-/nm [6] |

**Table S1. Parameters used for MC simulation.** Parameters used for Monte Carlo simulations to generate equilibrium DNA configurations.
